# Engineered superinfective Pf phage prevents dissemination of *Pseudomonas aeruginosa* in a mouse burn model

**DOI:** 10.1101/2022.12.05.519246

**Authors:** Federico I Prokopczuk, Hansol Im, Javier Campos-Gomez, Carlos J. Orihuela, Eriel Martinez

## Abstract

Pf is a filamentous bacteriophage integrated in the chromosome of most clinical isolates of *Pseudomonas aeruginosa*. Under stress conditions, mutations occurring in the Pf genome result in the emergence of super-infective variants of Pf (SI-Pf) that are capable of circumventing phage immunity; therefore SI-Pf can even infect Pf-lysogenized *P. aeruginosa*. Herein, we identified specific mutations located between the repressor and the excisionase genes that result in the emergence of SI-Pf. Based on these findings, we genetically engineered a SI-Pf (eSI-Pf) and tested it as a phage therapy tool for the treatment of life-threatening *P. aeruginosa* infection of burns caused by strain PAO1. eSI-Pf was able to infect PAO1 biofilms formed in vitro on polystyrene and inhibited their formation when at high concentration. eSI-Pf also infected PAO1 present in burned skin wounds on mice but was not capable of maintaining a sustained reduction in bacterial burden beyond 24 hours. Importantly, and despite not lowering CFU/g of burn skin tissue, eSI-Pf treatment completely abolished the capability of *P. aeruginosa* to disseminate from the burn site to internal organs. Over the course of 10 days, this resulted in bacterial clearance and survival of all treated mice. We determined that eSI-Pf induced a small colony variant of *P. aeruginosa* that was unable to disseminate systemically in our burned mouse model during acute infection. Our results suggest that eSI-Pf has potential as a phage therapy against highly recalcitrant antimicrobial resistant *P. aeruginosa* infections of burn wounds.

**IMPORTANCE:** *Pseudomonas aeruginosa* is a major cause of burn related infections. It is also the most likely bacterial infection to advance to sepsis and result in burn-linked death. Frequently, *P. aeruginosa* strains isolated from burn patients display a multidrug resistant phenotype necessitating the development of new therapeutic strategies and prophylactic treatments. In this context, phage therapy using lytic phages has demonstrated exciting potential in the control *P. aeruginosa* infection. However, lytic phages have a set of drawbacks during phage therapy including the induction of bacterial resistance and limited bacteria-phage interactions in vivo. Here we propose an alternative approach to interfere with *P. aeruginosa* pathogenesis in a burn infection model, i.e., using an engineered super-infective filamentous phage. Our study demonstrates that treatment with the engineered Pf phage can prevent sepsis and death in a burn mouse model.

## Introduction

*Pseudomonas aeruginosa* is a major cause of acute and chronic infections including those affecting the urinary tract, lung, and skin (1, 2). This opportunistic Gram-negative bacterium is in particular a major health problem in nosocomial settings as isolates are commonly multi-drug resistant. For this reason, the World Health Organization (WHO) has classified *P. aeruginosa* as a critical pathogen that urgently requires new therapeutic strategies (3). The low success rate and overall slowdown in the identification of new and efficacious antibacterial drugs underlies the pressing need to develop alternative interventions.

Bacteriophage are viruses that infect bacteria and, in some instances, cause their lysis (4). In Eastern Europe, bacteriophage have been used for over a century to treat recalcitrant bacterial infections. Recently, the potential of phage therapy has been explored in other countries and bacteriophage are increasingly being used as a last-ditch effort to treat lifethreatening infections caused by multi-drug resistant bacteria (5). Along such lines, the use of bacteriophages to treat advanced *P. aeruginosa* lung infection among those with cystic fibrosis has been conducted for years, and the results so far have been encouraging (6, 7).

Filamentous phage are characterized by their long and slender morphology which consists of a proteinaceous tube encasing a positive-sense single-stranded circular DNA genome (8). They belong to the *Inovirus* genus of the *Inoviridae* family of phages (9). Filamentous phages replicate their DNA using the rolling circle replication mechanism, a process of unidirectional nucleic acid replication that rapidly synthesizes multiple copies of the genome. In some instances, the genome of filamentous phage integrates into the bacterial chromosome and there remains as a lysogenic phage until specific conditions induce its excision and re-initiation of DNA replication, viral assembly, and in turn horizontal transmission of the phage (10). Importantly, bacteria lysogenized with a filamentous phage are typically resistant to infection with the same phage type as result of a mechanism called phage immunity (11, 12). The mechanism involves the expression of the phage repressor from the chromosome integrated copy which in turn binds to the DNA of the infecting phage and prevents its replication (12). Notably, and unlike most other phages, filamentous phages typically exit the bacterial cell without killing its host. This is because the phage is assembled and released from the bacterium’s membrane through a process known as extrusion (13). Accordingly, filamentous phages are powerful and popular molecular tools due to the fact that they can easily be genetically manipulated and, in turn, used to induce the expression of proteins by the target bacterium (14).

Pf is a filamentous bacteriophage which infects and can be found in the chromosome of most clinical isolates of *P. aeruginosa* (15); up to 68% of clinical strains of *P. aeruginosa* harbor a Pf phage (16). When grown in vitro, *P. aeruginosa* accumulate up to 10^11^ particles of Pf per milliliter of culture media. In vivo, Pf is also produced, and has been isolated from patients with cystic fibrosis at concentrations as high as 10^7^ particles per milliliter of sputum (17). Under stress conditions, *P. aeruginosa* produces a Pf variant that can circumvent phage immunity and successfully infects Pf-lysogenized bacteria (18). The exact mechanism leading to the emergence of these super-infective (SI) variants is unclear, but studies have suggested that stress-mediated mutations in the intergenic region between the phage repressor and the phage excisions are responsible(19). Notably SI-Pf infection can result in the lysis of *P. aeruginosa*. However, *P. aeruginosa* that survive SI-Pf infection are transiently resistant to Pf induced plaques and are deficient in twitching motility (20). The mechanism involves expression of the Pf phage protein PfsE termed super-infection exclusion factor (20).

Pertinent to our study, a recent report by Tortuel *et al*. showed that *P. aeruginosa* infected with SI-Pf phage had reduced virulence in lettuce plant and *Caenorhabditis elegans* infection models (21). Tortuel *et al*. reported decreased spread of the SI-Pf within the lettuce leaf, as well as a reduced rate of *C. elegans* mortality when compared to control, respectively. This observation led us to question whether SI-Pf has potential as a novel therapeutic strategy to treat humans with life-threatening infections of *P. aeruginosa*. In support of this notion, filamentous phages are thought to be safe for use in mammals (22); to our knowledge no record of adverse effects of Pf or bacteriophage on an animal or person has been reported. Yet filamentous phages have not been considered as primary tools for phage therapy since they do not generally lyse the infected bacterium and therefore would not eradicate the bacterium by direct means. In this study we created an engineered eSI-Pf phage and tested its potential to ameliorate *P. aeruginosa* infection of a major burn injury. Notably, treatment with eSI-Pf dramatically suppressed bacterial dissemination from the skin to internal organs and prevented mice from dying as compared to untreated controls.

## Results

### Identification of mutations resulting in super infective Pf phage

SI-Pf variants emerge spontaneously under stress conditions. This is due to mutations in the phage genome that result in excision from the bacterial chromosome with subsequent dysregulation of replication (23, 24). Along such lines, we have reported that a *P. aeruginosa* mutant lacking the mismatch repair enzyme MutS spontaneously produced SI-Pf at a high frequency (25). Transition of Pf to the SI-Pf state in this mutant was deemed to be due to the hypermutagenic phenotype caused by *mutS* disruption. Importantly, the specific mutations in the Pf chromosome that result in a super-infective phage were not well delineated. While some studies have detected mutations in the phage repressor and in the intergenic region between the repressor and the exsicionase gene, other have reported mutations in other phage genes such as PA0723, PA0724 and PA0725 (21). None of these had been experimentally validated as being at cause.

To determine the specific mutations responsible for the emergence of SI-Pf, we streaked out to individual colonies a version of *DmutS* that did not yet produce SI-Pf. Forty individual clones of *DmutS* in PAO1, which is already lysogenized with Pf, were cultured independently in tryptic soy broth (TSB) culture media and filtered supernatants were used to infect a PAO1 lawn to obtain plaques of diverse SI-Pf variants. A single plaque was isolated for each supernatant, respectively, and the phage variants from each plaque were expanded to isolate their replicative form (RF) for sequencing (**Fig. 1A**). In all the forty RFs corresponding to SI-Pf were sequenced. Mutations associated with the formation of the SI-Pf were all localized to the intergenic region between the Pf repressor and the excisionase gene (**Fig. 1B**). The frequency of each mutation is shown in **Figure 1B**. The most abundant mutant was A32G (62.5%), whereas the second most abundant mutation was A6G (17.5%). We therefore chose to introduce both mutations into the Pf genome in an attempt to recreate the SI phenotype. These mutations were introduced into the Pf genome using inverse PCR as described in the Methods section. Notably, replacement of A32 by G or A6 by G led to the creation of a phage variant that produced plaques on a lawn of lysogenized PAO1. The mutations in these areas are sufficient to confer the SI-Pf phenotype.

**Figure 1.**
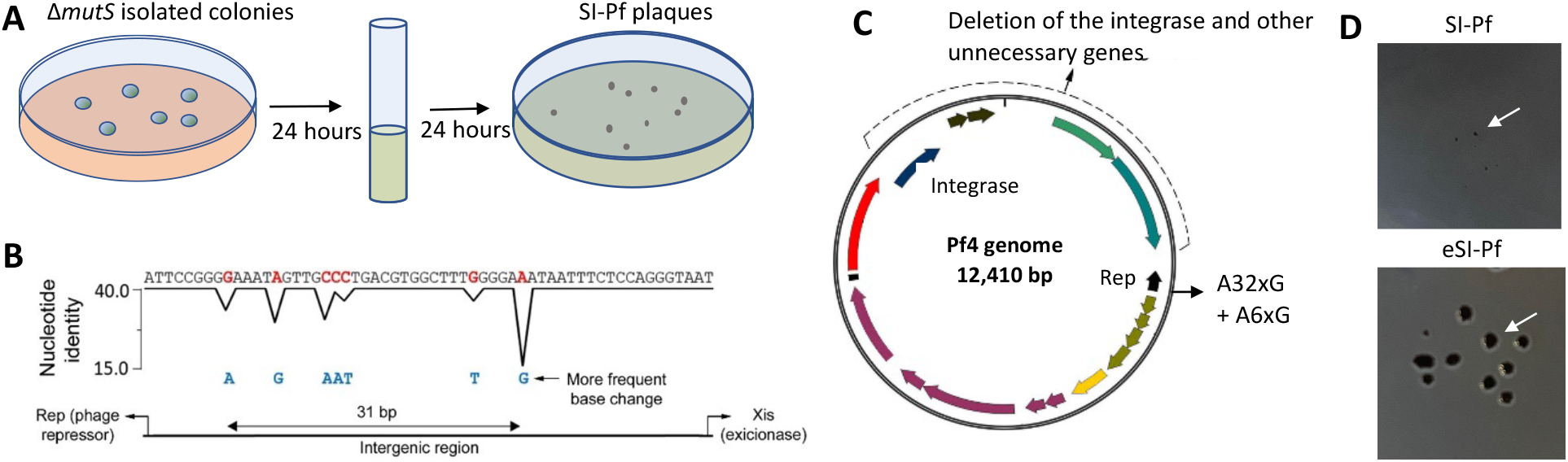
Construction of an engineered SI-Pf phage. **A)** Screen strategy to identify single mutations leading to the formation of the SI-Pf. Forty isolated colonies of the *mutS* mutant_were cultured in LB broth and the supernatant used to infect_a PAO1 lawn to select isolated plaques **B)** Sequence analysis of 40 SI-Pf individual clones isolated from single plaques. All mutations were localized in the intergenic region between the phage repressor and the excisionase genes. **C)** The most frequent mutations A32xG and A6xG were introduced in the replicative form of Pf4 phage. Additionally, all genes unnecessary for replication, assembly and export of Pf, including the integrase encoding gene, were deleted from the phage genome. **D)** Supernatant of *DmutS* (top) or PAO1 infected with eSI-Pf (bottom) were diluted to obtain phage isolate plaques on a PAO1 lawn. eSI-Pf formed larger plaques than Pf with a spontaneous SI-Pf phenotype. Images were obtained with a hand held camera at equal distance and magnification.

### Creation of eSI-PF

One of our goals was to test whether SI-Pf has utility as a potential therapeutic for recalcitrant *P. aeruginosa* infection. Accordingly, we created a version of eSI-Pf that in addition to the A32xG and A6xG mutations had a deletion encompassing the integration site (attP), the integrase, and other genes out of the phage core. This deletion was added so as to reduce the chances of eSI-Pf reversion to wildtype as well as to prevent its ability to integrate into the chromosome of the bacterium (**Fig. 1C**). When used to infect a PAO1 lawn, eSI-Pf formed larger plaques than spontaneously generated SI-Pf (Fig 1D).

### The engineered superinfected phage infects *P. aeruginosa* biofilm in vitro and in vivo

Pf is highly induced when *P. aeruginosa* is within a mature biofilm (26). Moreover, the phage is a component of the biofilm matrix of *P. aeruginosa* that carry the phage, suggesting that Pf may have special physicochemical characteristics that impact its interaction with the biofilm matrix. We considered that this feature might negatively influence the ability of exogenous eSI-Pf to infect *P. aeruginosa* in a biofilm. To test accessibility of eSI-Pf to *P. aeruginosa* within a biofilm we created an eSI-Pf version that encoded the tdTomato fluorescent protein, which results in eSI-Pf infected bacteria having red fluorescence. The successful expression of the TdTomato gene in PAO1 grown on tryptic soy blood agar plates infected with eSI-Pf-tdTomato was confirmed by fluorescent microscopy, which revealed a red ring around eSI-Pf-tdTomato plaques formed over a PAO1 lawn (**Fig. 2A**). Subsequently we tested the capability of eSI-Pf-tdTomato to infect a 24-hour biofilm of PAO1 formed on polystyrene plates. Representative images shown in **Figure 2B** demonstrate that a meaningful subset of PAO1 now expressed tdTomato (red), indicating they had been successfully infected with eSI-Pf-tdTomato. We enumerated this, and in a 24-hour biofilm that had be super infected with eSI-Pf-tdTomato for 24 hours, 30-35% of all bacteria detected were fluorescing red. Finally, and to demonstrate that eSI-Pf could also infect *P. aeruginosa* in vivo and under clinically relevant conditions, mice that had suffered burn injury and been infected with PAO1 were treated with eSI-Pf-tdTomato at the lesion site. Burned skin was collected after another 24h, sectioned, and analyzed by fluorescent microscopy. As shown in **figure 3C**, eSI-Pf-tdTomato successfully infected *P. aeruginosa* present at the infected wound site.

**Figure 2.**
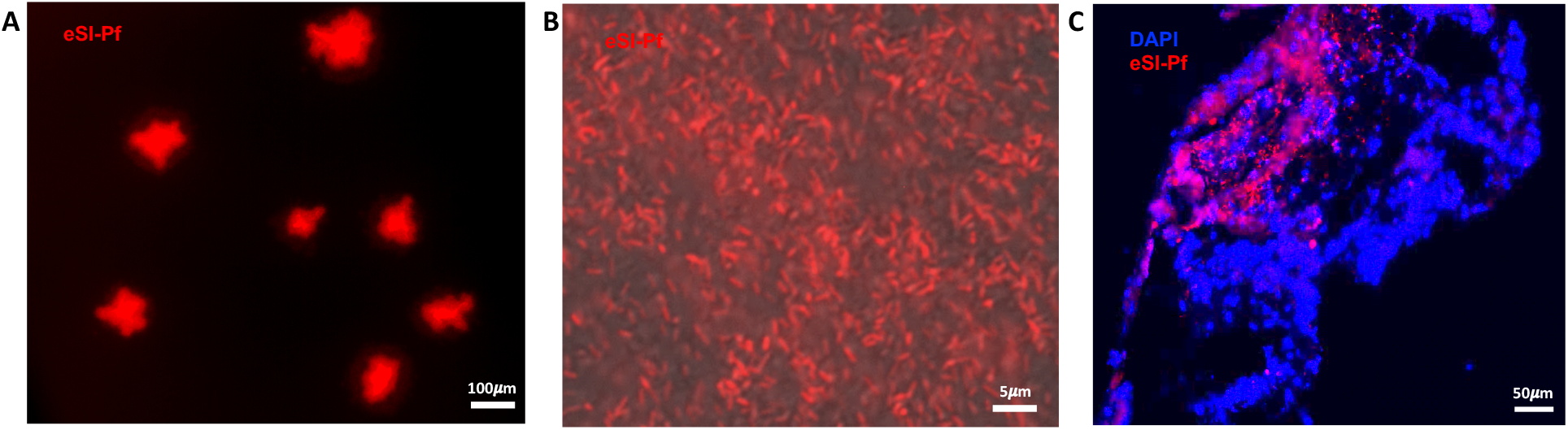
SI-Pf infects *P. aeruginosa* biofilm in vitro and in vivo. **A)** Fluorescent microscopy imagen of isolated plaques created by eSI-Pf-tdTomato on a PAO1 lawn. The scale bar represents 100*μ*m **B)** Fluorescent microscopy imagen of a 24 hours biofilm of PAO1 infected with eSI-Pf-tdTomato, red. The scale bar represents 5*μ*m **C)** Fluorescent microscopy imagen of a burn mice skin section infected with PAO1 for_24 hours and then treated intradermally with eSI-Pf-tdTomato. DAPI-stained nuclei, blue; PAO infected with eSI-Pf-tdTomato, red. The scale bar represents 50*μ*m.

**Figure 3.**
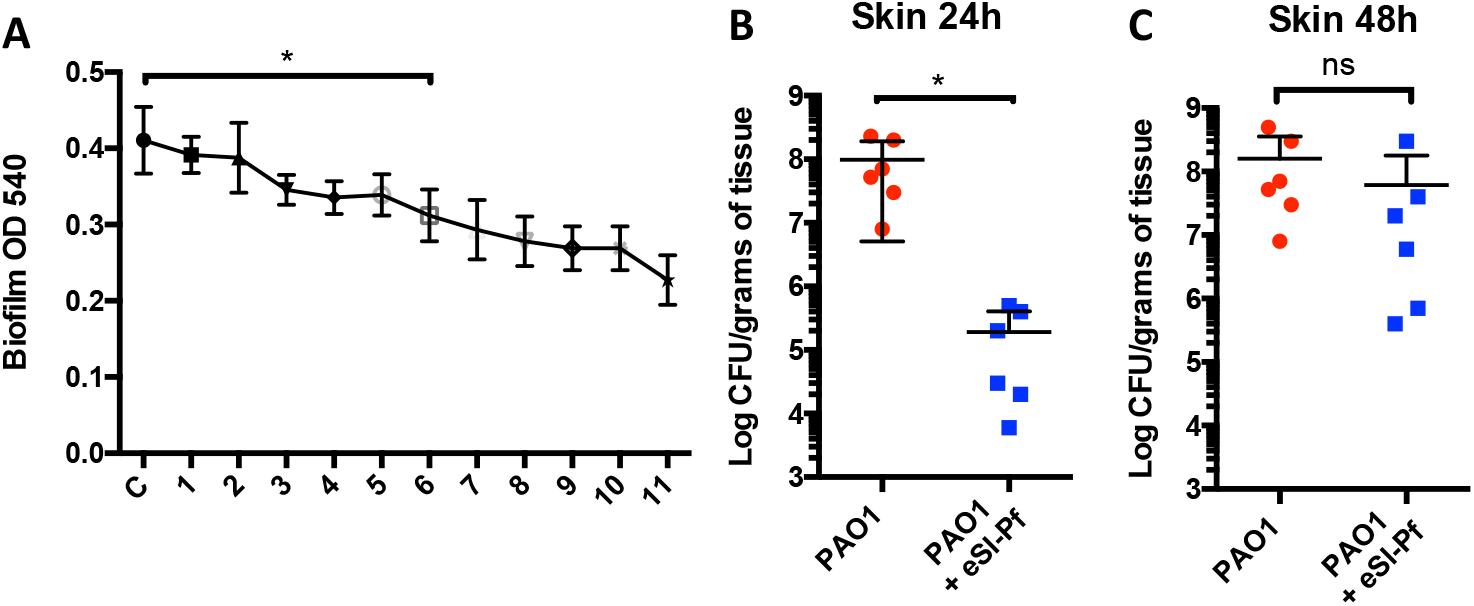
eSI-Pf disrupted stablished *P. aeruginosa* biofilms. **A)** A 24 hours biofilm of PAO1 was treated with eSI-Pf. Quantification of the biofilm was performed by crystal violet 4 hours post-treatment. Treatment with 10^6^ PFU/mL of eSI-Pf or higher concentration was detrimental for the biofilm. **B)** Mice infected for 24 hours with PAO1 were treated with eSI-Pf. 24 hours posttreatment the skin showed reduced bacterial load compared to control. **C)** However, no significant difference was detected 48 hours posttreatment. Statistics: one-way ANOVA with Tukey’s multiple comparison (Panel A) Student’s t test (panels B and C). Each dot is a biological replicate. Errors bars represent the standard error of the mean. Asterisks denote statistical significance. *, *P* < 0.05; ns, not significant.

### High titers of eSI-Pf temporally impact stablished biofilm *in vitro* and *in vivo*

Biofilm formation is a critical event in *P. aeruginosa* pathogenicity and the altered metabolism of bacteria growing in biofilm conditions provides intrinsic antimicrobial resistance (27). Biofilm formation also impairs host immune cell clearance and therefore slows wound healing and can lead to a chronic infection state (28). While production of the WT version of Pf has been described to promote biofilm formation by creating a liquid crystalline organization of the biofilm matrix (29), the consequence of superinfective phage, if any, on the stability of established biofilms was unclear. We observed that the addition of exogenous eSI-Pf to 24h biofilms formed in vitro was detrimental. This occurred in a concentration dependent manner with a clear effect observable when we added more than 10^6^ PFU/mL (**Fig. 3A**). However, this effect was limited, and the addition of eSI-Pf did not lead to the complete collapse of the biofilm structure even when treated with as much as 10^11^ PFU/mL. Importantly, we obtained comparable results with regard to bacterial eradication in vivo. One day after burn injury and infection with PAO1, mice were treated intradermally with eSI-Pf at the site of infection. When infected burned skin was isolated and *P. aeruginosa* quantified, mice treated with eSI-Pf showed significant reduction in bacterial load 24 hours post-treatment (**Fig. 3B**). However, the analysis of the skin samples at 48 hours treatment revealed that the bacterial load had recovered (**Fig. 3C**).

### eSI-Pf abolished bacterial dissemination in a mouse burn model

Despite not resulting in sustained reduction of bacterial burden at the burn site, we observed a drastic effect of eSI-Pf treatment on the ability of *P. aeruginosa* to disseminate from the skin to internal organs. In our burn model, mice infected with 10^6^ CFU of PAO1 had up to 5.0 x 10^4^ CFU/gram of tissue in the spleen (**Fig. 4A**) and similar levels in the liver (**Fig. 4B**). However, no bacteria were detected in the spleen or liver of infected mice that had been treated with eSI-Pf 24h after initial bacterial challenge (**Fig. 4AB**). This was also stark evidence of protection in survival experiments performed in parallel. All eSI-Pf treated mice survived to the infection (**Fig. 4C**). Notably, treated mice recovered alertness and were responsive to interaction in contrast to untreated controls (**Video S1**). In addition, eSI-Pf treated mice were eventually able to eradicate the bacteria from the skin (ten days post infection), whereas all the untreated animals died in the first 4 days postinfection.

**Figure 4.**
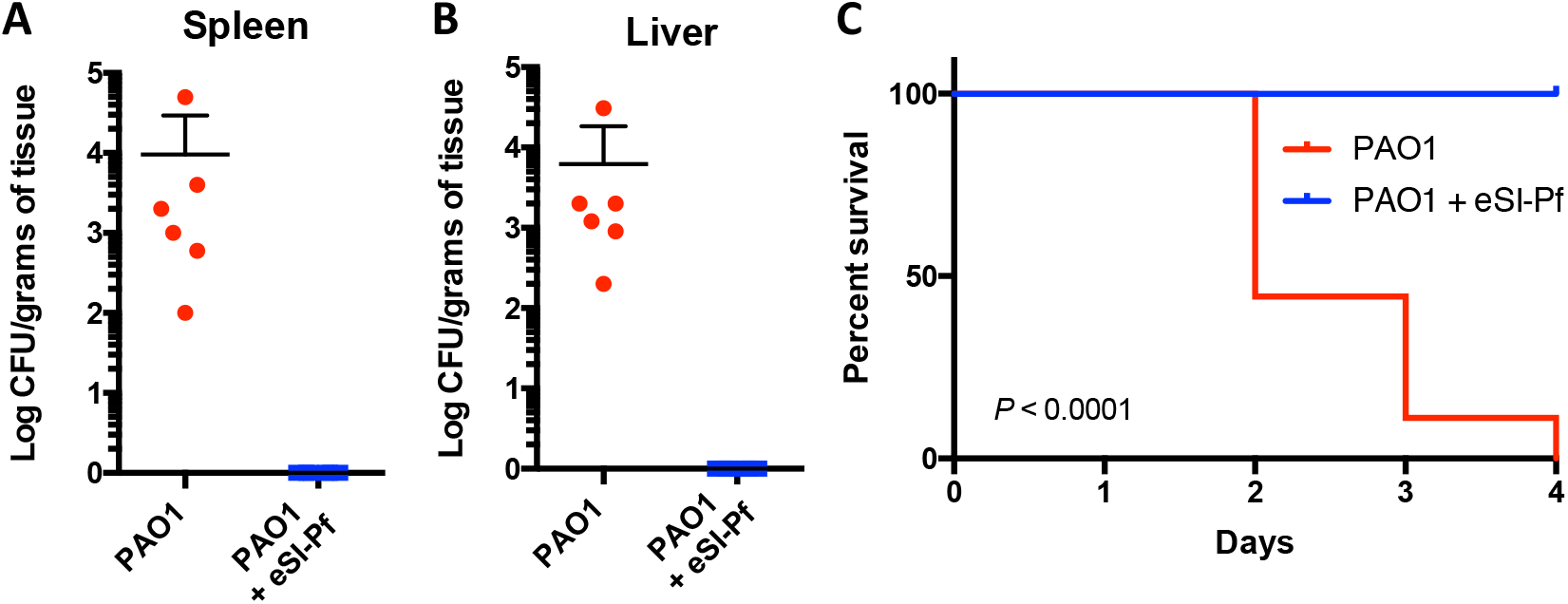
eSI-Pf leads to decreased *P. aeruginosa* virulence in a mouse burn model. **A)** PAO1 infected with eSI-Pf showed impaired dissemination to the spleen and **B)** to the liver. **C)** Survival curve of mice treated eSI-Pf and untreated control. Mice treated with eSI-Pf survived the infection. Statistics: Student’s t test (panels A and B). Mantel-Cox test (Panel C). Each dot is a biological replicate. Errors bars represent the standard error of the mean.

Since eSI-Pf treatment did not maintain a sustained reduction in bacterial load, we considered the possibility that *P. aeruginosa* that survived eSI-Pf challenge became attenuated and were therefore unable to disseminate. To test this, we treated PAO1 with eSI-Pf in vitro. Notably, infection of *P. aeruginosa* with SI-Pf resulted in formation of small colony variants (SCVs) that were resistant to the phage on solid medium (**Fig. S1A**)(18). Importantly, the proportion of SCVs obtained in vitro by treatment with eSI-Pf was proportional with the phage concentration (**Fig. 5A**). In agreement with previous reports, isolated SCVs showed dysregulated pigment production (**Fig. S1B**) and impaired motility (**Fig. S1C**) (20). We subsequently isolated 6 independent SCVs and treated burn-injury mice with 10^6^ CFU/mL each. Mice infected with SCVs were able to successfully colonize the skin of mice although less efficiently than WT PAO1 (**Fig. 5B**). In stark contrast to WT, we did not detect dissemination to internal organs of mice treated with SCVs. (**Fig. 5D**). What is more, all mice treated with SCV survived the infection (**Fig. 5E**).

**Figure 5.**
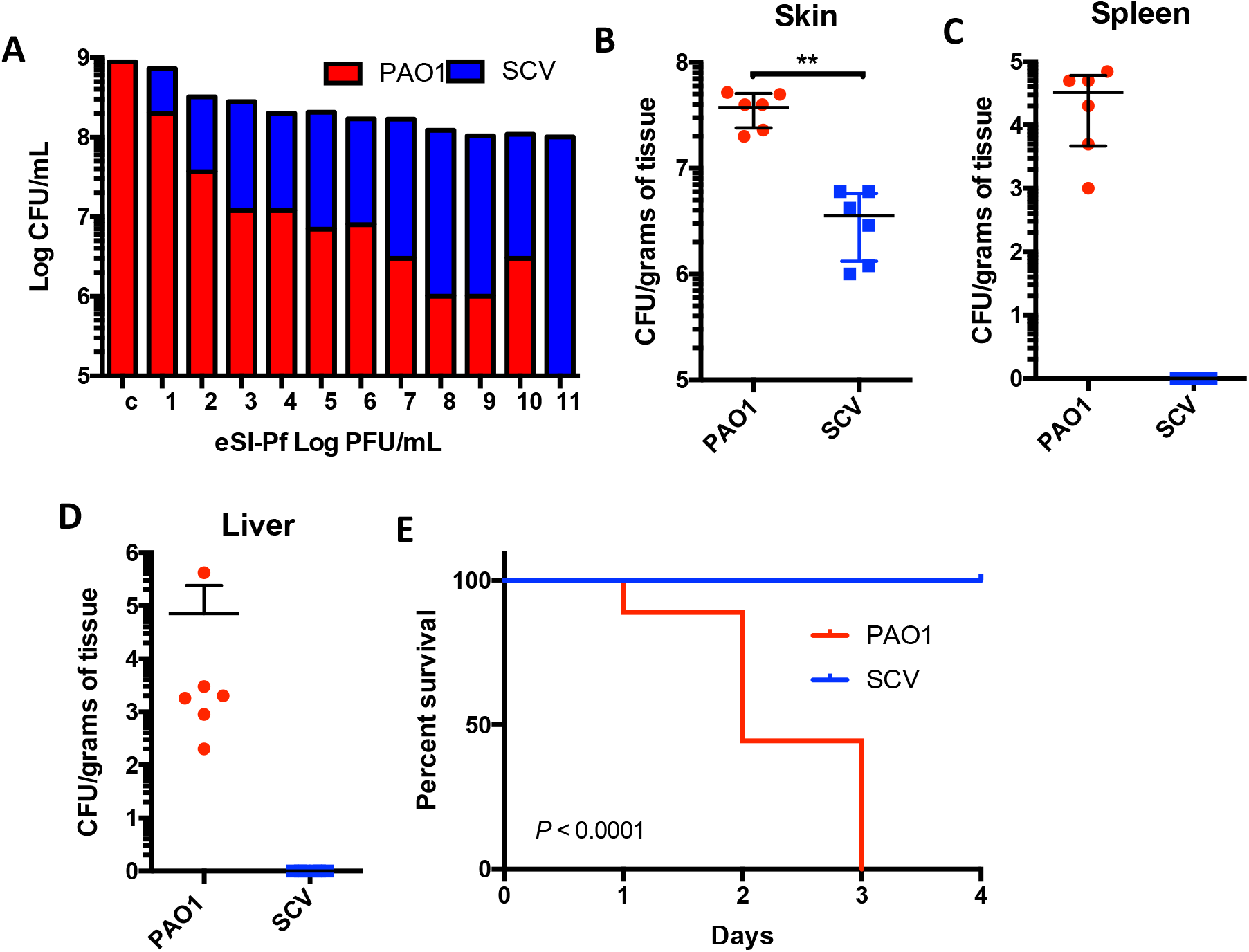
eSI-Pf infection of *P. aeruginosa* resulted in an attenuated phenotype in a mouse burn model. **A)** eSI-Pf infection of *P. aeruginosa* induces formation of small colony variants (SCV). This occurred in a phage concentration dependent manner. **B)** Burn mice were infected with SCVs. The SCV were able to colonize the burn skin but less efficient that PAO1 WT. **C)** The SCV showed impaired dissemination to the spleen and **D)** to the liver **E)** Survival curve of mice infected with the SCV versus PAO1 WT. The SCVs showed an_attenuated phenotype. Isolated SCVs Statistics: Student’s t test (panels B, C and D). Mantel-Cox test, P < 0.0001 (Panel E). Each dot is a biological replicate. Errors bars represent the standard error of the mean. **, *P* < 001.

## Discussion

*P. aeruginosa* is the most prevalent bacteria major burn wound infections (30). It is also the most common bacterial burn infection to advance to sepsis and lead to death. During burn-site infection *P. aeruginosa* behaves more aggressively when compared to other types of wound or airway infections (31). In these latter instances, *P. aeruginosa* infects without immediate systemic spread or mortality (32). This may be due to the known immunosuppressive effects of large burns. Remarkably, up to 58% of clinical samples from wounded patients are multidrug resistant, suggesting that innovative therapeutic and prophylactic treatments are urgently needed (33).

Phage therapy has been in use in Eastern Europe for almost a century as a means to treat bacterial infection. However, the until recent efficacy of antimicrobials has eclipsed phage therapy research. Now, in an era of worldwide dissemination of antimicrobial resistance and with the current capabilities to molecularly identify and characterize phages, their use as a treatment modality is an increasingly attractive option for recalcitrant bacterial infections (34). Other advantages of phages as therapeutic tools include that they are a self-replicating, which makes their production fast and affordable compared to antibiotics. Phage also have specificity for their bacterial hosts and do not cause allergic reactions (22). Pertinently, phage therapy has been demonstrated to have efficacy in the control *P. aeruginosa*, principally among cystic fibrosis patients who have lung infections with multidrug resistant strains (34). However, these therapies are based on the use of overtly lytic phages, such as PB1-like, phiKZ-like, and LUZ24-like phages and present a distinct set of challenges such as emergence of bacterial resistance to the phage and limited bacteria-phage interactions in vivo (35).

Herein, we developed an engineered SI-Pf phage to treat *P. aeruginosa* infections in a burn context. Our thought being that SI-Pf phage has the ability to overcome pre-existing phage immunity and therefore had potential to be used to treat infection, even those caused by Pf lysogenized strains. One strength was the demonstrated ability of Pf to infect *P aeruginosa* within a biofilm (29). The rationale for our study were the prior observations by Tortuel *et al*. revealing *P. aeruginosa* infected with SI-Pf has reduced virulence in lettuce and *C. elegans* infection models (21). These models, have demonstrated value in identifying *P. aeruginosa* virulence factors and therefore hinted that SI-Pf treatment of infected wounds may ameliorate infection in mammals (36). Our own results using a pre-clinical burn mouse model showed this was the case as eSI-Pf infected PAO1 were unable to disseminate from the injury site to the spleen and liver, and all treated animals eventually cleared the infection and survived challenge. In contrast, all burned mice that did were not the recipient of eSI-Pf phage therapy died, highlighting the severity of this type of infection and the importance of having alternatives to antimicrobials readily available should they fail.

Use of SI-Pf to reduce virulence opens a new possibility for a combinatorial strategy, and importantly does not exclude the complementary use of antimicrobials. For example, previous studies have observed increased sensitivity to antimicrobials upon filamentous phage infection (37). Specifically, Pf infecting LESB58 strain had increased susceptibility to ciprofloxacin, gentamicin, ceftazidime, and streptomycin while Pf infection PA14 showed increased sensitivity to ciprofloxacin and ceftazidime (24). This effect of Pf in re-sensitizing newly infected strains to antibiotics have itself a great therapeutic implication. We speculate that the explanation for this includes an altered metabolic need of the bacterium when forced to accommodate phage replication. Again, further studies are required to determine the impact of SI-Pf on antimicrobial susceptibility in the context of an infection setting and whether this enhances killing or resistance. Studies that have used phage as a therapeutic lytic treatment concluded that phage cocktail is more effective than a single phage (38). In the same vein, a combinatorial treatment of phages and antibiotics is likely even more effective (39). For example, bacteriophage cocktail-antibiotic combination therapy against extensively drug-resistant *P. aeruginosa* infection allowed liver transplantation in a toddler with recalcitrant infection (40). The rationale is that the target for this phage may be selected against, and therefore multiple venues of attack make evolutionary escape less likely.

*P. aeruginosa* is undergoes mutation during infection with selection of favored mutants by the host environment (41). An eminent example of this is the emergence of SI-Pf during stress conditions. Along such lines, small colony variants (SCV) are also a common emergent phenotype frequently isolated from chronic infection sites (42). For *P. aeruginosa* the SCV phenotype is caused by genetic mutations or large scale inversions that often, but not exclusively lead to elevated intracellular levels of *c*-di-GMP(43). Although, SCV are frequently isolated from infections sites or in vitro conditions that favor constitutive biofilm formation, such as chronic infection and antibiotic treatments, other stress conditions can also lead to SCV formation in *P. aeruginosa* (44). For example, investigators have shown that *P. aeruginosa* in planktonic culture exposed to SI-Pf results in the emergence of SCV (18). In our study, and using our engineered SI-Pf variants, we reproduced this observation. Importantly, and while SCV emergence may promote low-grade chronic infection by emergence of an escape form, very little is known with regard to the impact of SCV formation during acute infection such as which occurs at burn sites. Here, using our mouse burn model we showed that SCV derived from treatment with eSI-Pf display an attenuate phenotype and are defective in dissemination from the infection site to internal organs. Determination of the exact molecular reason for SCV formation is an area of current research in our laboratory.

Regardless of the promising results obtained in our study, the use of the SI-Pf as a therapeutic strategy should be approached with considerable caution. Several studies have revealed that production of WT Pf promote *P. aeruginosa* pathogenicity in several ways. First, Pf serves as a structural element in *P. aeruginosa* biofilms (29, 45). In addition, Pf can trigger maladaptive anti-viral immune response that can promote *P. aeruginosa* chronic infections (46). This however is with the common forms of Pf, whereas much less is known about the role of the super-infective variants eventually produced in the bacterial population. It has been suggested that extracellular DNA released by the eventual lysis caused by Pf superinfection can be used as a structural component of the biofilm matrix, but this need further confirmation (17). Independently, SI-Pf treatment benefit would depend on the ability of *P. aeruginosa* to control phage spread that would cause collapse of the entire population. In summary, this study increases our understanding of SI-Pf dynamics in a clinically relevant setting and evaluates a proof of concept that a modified SI-Pf phage can be used to interfere with *P. aeruginosa* pathogenesis in burn-related infections, further studies are warranted to determine if this is indeed the case.

## Methods

### Strains and culture conditions

*P. aeruginosa* strain PAO1 (obtained from the Manoil Lab at the University of Washington, Seattle, WA, USA) (47) and its isogenic mutant *ΔmutS* (25) were used in this study. The strains were routinely grown in LB medium, or LB containing agar when solid medium was required. Swimming, swarming and twitching motilities were studied using the methods described by Rashid and Kornberg (48).

### DNA isolation and manipulation

Plasmid DNA was prepared using a QIAprep Spin Miniprep kit (Qiagen), which was used also to purify the replicative form of Pf phage. Q5 polymerase and modification enzymes were used according to the manufacturer’s instructions (NE Biolabs). The DNA was visualized with ethidium bromide (1 μg/ml). The replicative forms of the SI-Pf variants were used for sequencing using new generation sequencing (NGS). The sequences were analyzed using SnapGene 5.2.4.

### Production and isolation of SI-Pf variants

A *mutS*-deficient strain which spontaneously produces superinfective Pf (SI-Pf) viral particles was used as the SI-Pf donor. Isolated colonies of *ΔmutS* were grown in 5 ml of LB for 20 h with shaking (240 rpm) at 37°C. One milliliter of the culture was spun down, and the supernatant was serially filtered through 0.45- and 0.22-μm-pore-size filters (sartorius). Cell-free supernatants containing SI-Pf particles were used to produce isolate phage plaques on a lawn of PAO1 lawn.

### Construction and purification of the engineered SI-Pf phage

Site-direct mutagenesis in Pf phage and fragment deletions were performed by inverse PCR using the replicative from of Pf phage and the primers; Pf-RV-NotI: GACAGCGGCCGCGTCAGCTTCATTTCTTGCCTTCCA and Pf-FW-NotI: GACAGCGGCCGCCAGGTCCTTTGCCACCAACC. For site-direct mutagenesis, A32G was created using primers with mutagenic ends; A32G-FW: GATAATTTCTCCAGGGTAATTATTTCTCTAGC; A32G-RV: TCCCCAAAGCCACGTCAG; A6G-FW: GGTTGCCCTGACGTGGC and A6G-RV: ATTTCCCCGGAATAAATTTCTATATGAGCAC. To create a SI-Pf version expressing tdTomato we digested PUC-tdTomato vector with notI and the fragment containing the tdTomato gene was cloned into the replicative form of the eSI-Pf. The supernatant of transformed PAO1 was analyzed for red plaque formation a PAO lawn. To purify eSI-Pf, the supernatant of PAO1 infected with the eSI-Pf was serially filtered through 0.45- and 0.22-μm-pore-size filters (sartorius). Phage particles were precipitated from the filtered supernatant by adding sodium chloride and polyethylene glycol to give final concentrations of 3 and 5% (wt/vol), respectively. The mixture was incubated on ice for 30 min and centrifuged at 12,000 × *g* for 20 min. The supernatant was discarded, and the phage-containing pellet was resuspended in 1 ml of phosphate-buffered saline.

### Phage titration

Titers of SI-Pf were routinely determined by dropping serial dilutions of filtered culture supernatants of phage-producing cells or purified particles onto soft LB agar (0.4%) plates containing the PAO1 strain. The numbers of plaques were determined after overnight incubation at 37°C. Isolated plaques were used to isolate monoclonal SI-Pf when required.

### Quantification of biofilm formation

Biofilm assays were performed following the O’Toole protocol (49). Briefly, *P. aeruginosa* strains were cultured overnight in LB agar plates at 37 °C. Bacterial suspensions were prepared in M63 medium to an OD_600_=1. Ten microliters of bacterial suspension were inoculated into each well of a 96-well microtiter plate containing 200 μl of M63 complete media. Biofilms were allowed to form at 30 °C overnight. For biofilm quantification the wells were washed two times with PBS and 200 μl of 0.1% crystal violet was added to the wells and incubated for 10 min. Wells were washed three times with phosphate-buffered saline, then crystal violet-stained biofilm was solubilized with 250 μl of 30% acetic acid and the absorbance was measured at 550 nm.

### Fluorescence microscopy

Visualization of the biofilm and phage plaques were conducted using Leica LMD 6 and Nikon eclipse Ti microscope. To make the biofilm structure, we directly used slide glass as surface and made the biofilm for 48 hours incubation. Phage was inoculated at the 24 hours timepoint. The fluorescence intensity was compared using red fluorescence (tdTomate-SI-Pf). For the plaque images, overnight culture of SI-Pf on PAO1 was used.

### Back burn mice infection

Mice infection was performed as previously described (50). Briefly, five-to six-week-old BALBc mice (Jackson Labs) were anesthetized with a mixture of xylazine and ketamine. The back was shaved with an electric clipper and then depilated with depilatory cream. The burn was induced using a hot bar and the infection was performed intradermally with 100 μl of bacterial suspension containing 10^6^ CFU of *P. aeruginosa* PAO1 or the small colony variant (SCV). The bacterial burden in the skin, liver and spleen was assessed by homogenizing the organ and plating dilutions on LB agar plates.

### Treatment of burn wound infection with eSI-Pf

Four hours post-infected mice were treated intradermally with 100μl of purified eSI-Pf containing 10^7^ PFU of eSI-Pf or 100 μl of phosphate buffer saline for controls. After treatment mice were monitored each day for symptoms and deaths for 10 days or euthanized 24 or 48 hours postinfection to determine bacterial burden in the skin, liver and spleen.

### Ethics statement

The animal experimental design was approved by the Institutional Animal Care and Use Committee at The University of Alabama at Birmingham, UAB (protocol no. IACUC-22197).

### Statistical analysis

Kaplan-Meier plots of mice survival experiments were compared using the log-rank (Mantel-Cox) test. We used 6 mice per condition. In the rest of the experiments oneway ANOVA or unpaired *t*-test was used. All statistical analyses were performed using GraphPad Prism 8 software (GraphPad Software, La Jolla, CA).

## Acknowledgments

This work was supported by NIH National Institute of Allergy and Infectious Diseases (NIAID) grants: 1R01AI172796-01, 5R01AI114800-08, 5R01AI156898-03, 5R21AI148368-02.

---

**Figure S1:**
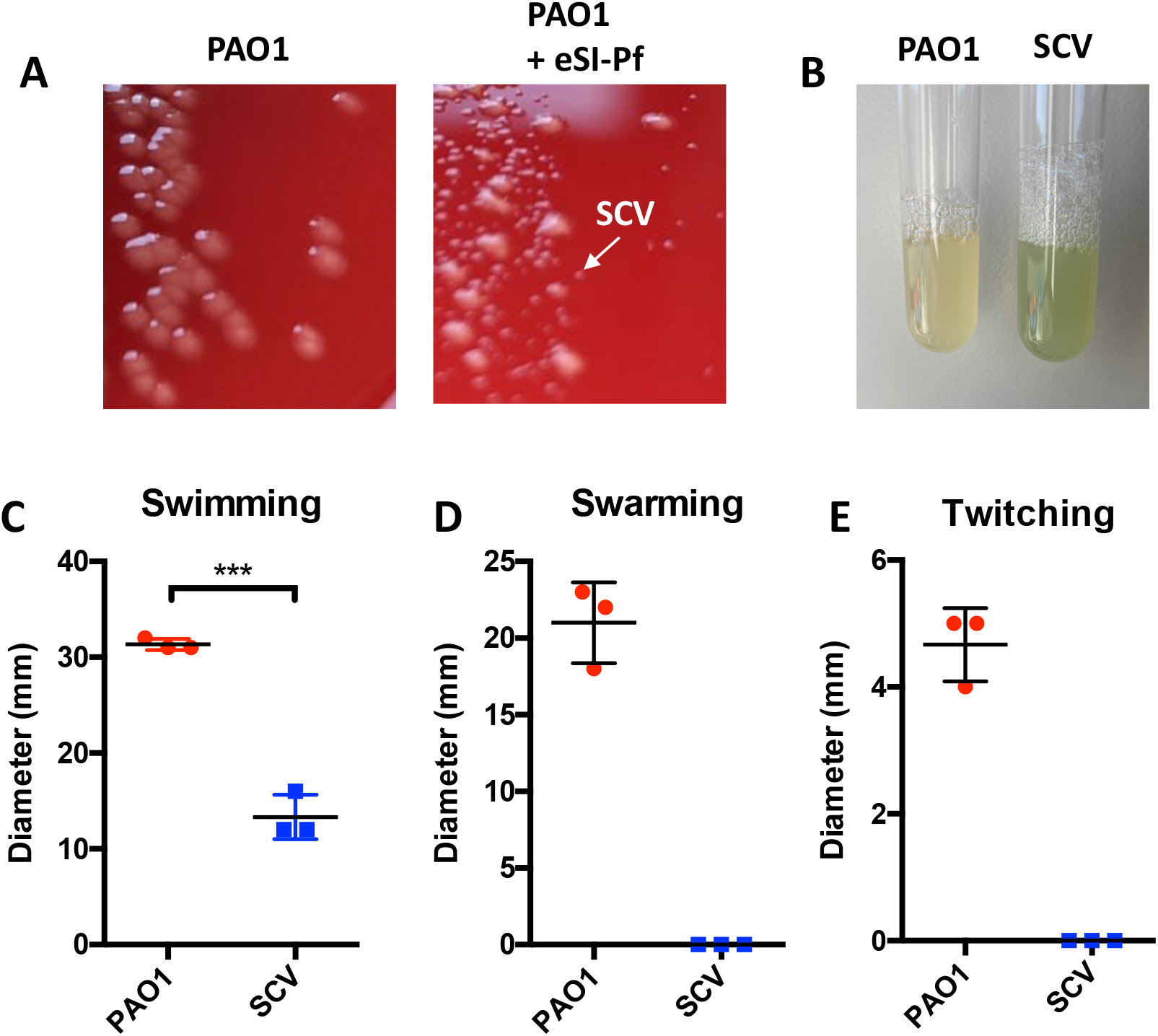
eSI-Pf infection induce small colony formation. **A)** Left: untreated PAO1, Right: eSI-Pf treated PAO1. The arrow indicates a representative small colony variant (SCV). **B)** PAO1 and a SCV cultured overnight in TSB. The SCV shows dysregulation of pigment production. **C)** SCVs showed reduced swimming motility compared to the PAO1 WT phenotype. **D)** The SCVs showed impaired swarming and **E)** twitching motilities. Statistics: Student’s t test (panels C-E). ***, *P* < 0.001

**Video S1:** eSI-Pf treated mice are more alert and responsive than controls. Burned mice were infected with PAO1. Four hours post-infection, a cohort of mice were treated intradermally with eSI-Pf at the burn/infection site. Videos were taken at 24 hours postinfection Top: treated with eSI-Pf. Bottom: untreated control.

